# Chemotherapy Signatures Map Evolution of Therapy-Related Myeloid Neoplasms

**DOI:** 10.1101/2022.04.26.489507

**Authors:** Benjamin Diamond, Bachisio Ziccheddu, Kylee Maclachlan, Justin Taylor, Eileen Boyle, Juan Arrango Ossa, Jacob Jahn, Maurizio Affer, Tulasigeri M. Totiger, David Coffey, Justin Watts, Sydney X Lu, Niccolò Bolli, Kelly Bolton, Jae H. Park, Heather Landau, Karuna Ganesh, Andrew McPherson, Mikkael A. Sekeres, Alexander Lesokhin, David Chung, Yanming Zhang, Caleb Ho, Mikhail Roshal, Jeffrey Tyner, Stephen Nimer, Elli Papaemmanuil, Saad Usmani, Gareth Morgan, Ola Landgren, Francesco Maura

## Abstract

Patients treated with cytotoxic therapies, including autologous stem cell transplantation, are at risk for developing therapy-related myeloid neoplasms^1, 2^. Pre-leukemic clones (i.e., clonal hematopoiesis) are detectable years before the development of these aggressive malignancies^3-5^, though the genomic events leading to transformation and expansion are not well-defined. Here, leveraging distinctive chemotherapy-associated mutational signatures^6-12^ from whole-genome sequencing data and targeted sequencing of pre-chemotherapy samples, we reconstruct the evolutionary life-history of 39 therapy-related myeloid malignancies. A dichotomy is revealed, in which neoplasms with evidence of chemotherapy-induced mutagenesis from platinum and melphalan are relatively hypermutated and enriched for complex structural variants (i.e., chromothripsis), while neoplasms with alternative exposures bear a similar profile to *de novo* acute myeloid leukemia. Using chemotherapy-associated mutational signatures as a temporal barcode in each patient’s life, we estimate that several complex events and genomic drivers are acquired after chemotherapy exposure. In the case of treatment with high-dose melphalan and autologous stem cell transplantation, we demonstrate that the procedure allows clonal hematopoiesis to escape chemotherapy exposure entirely, and to be reinfused to expand to malignancy. This information reveals a novel mode of malignant progression for therapy-related malignancies that is not reliant on direct mutagenesis or even exposure to chemotherapy, itself, and prompts further investigation into leukemia-permissive effects of cytotoxic drugs.

---

Therapy-related myeloid neoplasms (tMN) have dismal a prognosis and their incidence is predicted to increase as cancer survival rates rise^13-15^. Some patients with underlying clonal hematopoiesis (CH) have a particularly high risk of tMN, especially those undergoing autologous stem cell transplantation (ASCT), a treatment with ubiquitous usage in lymphoproliferative disorders^3-5, 16^. It has been suggested that anti-cancer therapies exert positive selective pressure on CH bearing pre-leukemic driver mutations and that expansion is not contingent on the increased mutational burden relative to *de novo* AML^17, 18^ with speculation that other cellular machinery may be altered by exposure. Despite this, distinct DNA-damaging cytotoxic agents can measurably alter the mutational profile in each exposed cell, including in normal tissue^6, 19, 20^. The application of mutational signatures to whole genome sequencing (WGS) data has allowed for direct quantitation of chemotherapy-mediated DNA damage and revealed that a subset of tMN with prior platinum exposure do indeed have an increased, chemotherapy-specific, mutational burden as compared to non-platinum-exposed tMN^12^.

Chemotherapy-related mutational signatures are only detectable in bulk WGS following the clonal expansion of a single cell bearing its unique catalogue of chemotherapy-induced mutations (i.e., the single-cell expansion model, **Fig 1a**). The resultant mutational signature thus serves as a genomic single-cell barcode, linked to a discrete clinical and temporal exposure^6-9, 11, 21^. Here, we leverage chemotherapy-induced mutational signatures to measure genomic evolution of tMN with reference to each patient’s known therapeutic history. We focused the investigation on neoplasms that emerged following high-dose melphalan with ASCT because the chemotherapy is given in a single bolus, is associated with a distinct mutational signature (SBS-MM1)^8^, and because the leukapheresis procedure potentially allows for pre-leukemic clones to evade exposure to chemotherapy, altogether (**Fig 1b**). Overall, our data reveals two modes of expansion for tMN: one reliant on direct mutagenesis from chemotherapy to accrue late complex genomic events and the other dependent neither on direct chemotherapy-induced mutagenesis nor even direct exposure to cytotoxic agents.

## The mutational landscape of therapy-related malignancies

To measure the direct mutagenic activity of different chemotherapies and their roles in promoting tMN, we assembled a cohort of 40 tMN WGS from 39 patients with malignancies secondary to cytotoxic therapy (and/or radiation). Sixteen (40%) developed a secondary malignancy post-melphalan/ASCT (**Fig 1c, Extended Data Fig 1a, Extended Data Tables 1-3, Methods**). Latency between the diagnosis of primary malignancy and tMN varied (median, 5.5 years; IQR, 2.4-7.2 years; **Fig 1c**), and outcomes post tMN diagnosis were expectedly poor (**Extended Data Fig 1b**). As comparators, WGS of 21 *de novo* AML were imported from the TCGA processed with the same pipeline^22^ (**Methods**). To further gauge the effects of chemotherapy on genomic evolution of secondary malignancies, we sequenced five patients with B-ALL diagnosed post-melphalan/ASCT, two patients with multiple myeloma post-platinum, and one patient with transitional cell carcinoma post-melphalan/ASCT.

**Fig. 1.**
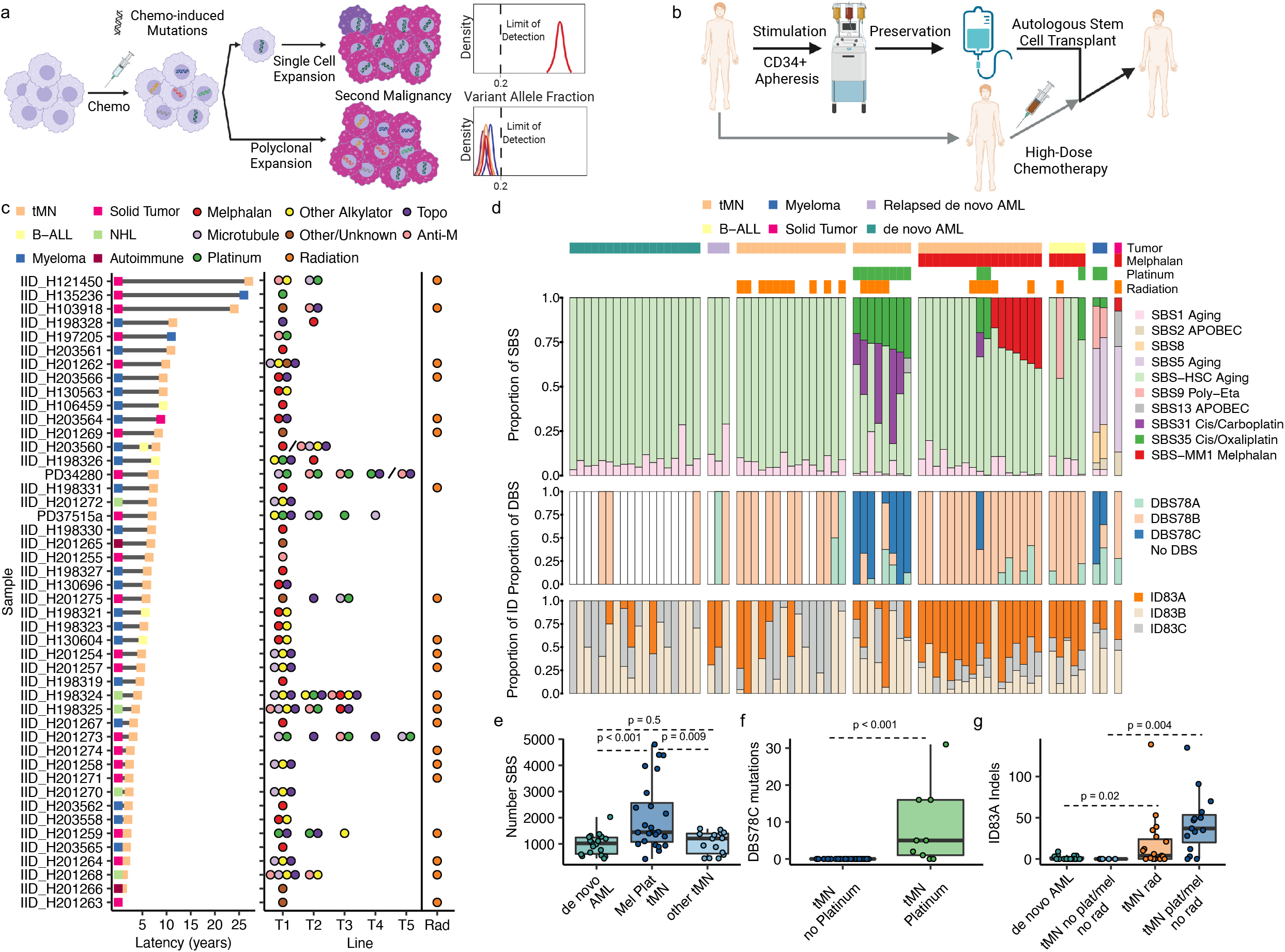
Genomic impact of chemotherapy on therapy-related tumors. **a)** Cartoon summarizing the single cell expansion model post-chemotherapy and its relevance in term of single nucleotide variants detectability. **b)** Cartoon depicting the procedure for autologous stem cell transplantation (ASCT). **c)** Therapy-related malignancies for which whole genome sequencing was performed with latency between primary tumor and second malignancy (left) and specific chemotherapy exposure (right). Backslashes separate sequential samples from the same patient. Anti-M, Antimetabolite. **d)** The proportional contribution of single base substitution (SBS), double base substitution (DBS), and indel (ID) mutational signatures in each tumor sample. Each column represents a unique patient. Samples are annotated by disease histology and therapy exposure. **e)** Boxplots for number of SBS for platinum/melphalan-exposed tMN as compared to *de novo* AML and other tMN. **f)** Boxplot for DBS among platinum-exposed and -unexposed tMN. **g)** Boxplot for indels signature mutations in *de novo* AML, tMN with radiation exposure, tMN without radiation or melphalan/platinum exposure, and tMN without radiation but with melphalan or platinum exposure. For e-f the p-values were estimated using Wilcoxon test.

We first compiled a list of high-confidence single-nucleotide variants (SNV) for each tumor and then performed mutational signature analysis following the previously published workflow based on *de* novo extraction, assignment, and fitting to identify and quantify the mutational processes that had been active in each tumor (**Methods, Fig1d, Extended Data Fig. 2, Extended Data Tables 4-7**)^8, 23^. Five known single-base substitution (SBS) mutational processes were identified in myeloid neoplasms: SBS1 and AML-HSC, attributable to intrinsic, clock-like mutations that accumulate with age, observed in all hematopoietic cells^12^; SBS31 and SBS35, attributable to mutations induced by intercalating platinum chemotherapies ^6^; and SBS-MM1, attributable to the alkylator melphalan^8, 11, 21^. As expected, *de novo* AMLs and their relapsed samples bore only evidence of clock-like mutational processes as neither of the induction agents cytarabine or anthracyclines are linked to distinct mutational signatures^7, 24, 25^. Overall, the only anti-neoplastic agents that induced measurable SBS mutagenesis were platinum and melphalan, with the mutational burden in tMN having exposure to these agents being significantly higher than in either *de novo* AML (Wilcoxon test, p<0.001) or unexposed tMN (Wilcoxon test, p = 0.009; **Fig 1e**). Though 5-Fluorouracil is known to leave a distinct SBS signature (SBS17a), no evidence of it was seen in this cohort, in line with its mechanism of mutagenicity on dividing – and not quiescent – cells^12^. Conversely, adducts generated by melphalan-and platinum-based agents are not dependent on cell turnover ^9, 12^ and induce mutagenesis in tumor and normal tissues^19, 20^. In fact, the mutational burden between *de novo* AML and tMN without melphalan or platinum exposure were strikingly similar (Wilcoxon test, p = 0.587; **Fig 1e**), adding evidence that chemotherapies typically associated with tMN may facilitate malignancy in absence of any appreciable increase in SBS mutagenesis^12^.

To further characterize the genomic impact of chemotherapy, we extracted double-base substitution (DBS) and indel (ID) signatures (**Methods, Fig 1d, Extended Data Fig 3**,**4, Extended Data Tables 4-5**,**8-9**). Mutations attributable to DBS78C, a signature resembling a combination of DBS4 and the platinum-associated E-DBS3^6^ (cosine similarity, 0.94), were identified in eight of ten tMN (80%) with prior platinum exposure as compared to none in unexposed samples (Wilcoxon test, p<0.001, **Fig 1f**). Three ID signatures were extracted, with ID83A being deconvoluted to the COSMICv3.2 ID8, previously linked to double strand breaks and ionizing radiation^7, 24^. ID83A was indeed enriched in radiation exposed tMN compared to unexposed *de* novo AML (Wilcoxon test, p=0.02), but also in tMN from patients with prior exposure to melphalan and platinum (Wilcoxon test, p = 0.004; **Fig 1g**) suggesting a link between this ID signature and therapy-mediated genotoxic damage^24^.

## Escape from chemotherapy-induced mutagenesis

All tMN with prior platinum exposure (n=10) had evidence of platinum-associated SBS31 and/or SBS35 signatures including samples with a latency from primary diagnosis (and exposure) until secondary malignancy up to 25 years (IQR, 3.9-9.0; **fig 1d**; **Extended Data Table 2**,**7**). This complete penetrance indicates that an originating cell, with direct DNA damage from platinum chemotherapy, expanded to clonal dominance (i.e., in line with the single-cell expansion model, **Fig 1a, Extended Data Fig 5a**)^11^. In striking contrast to the observations following platinum exposure, only 7 of 17 (41%) tMN with prior melphalan exposure had the SBS-MM1 melphalan signature, with neoplasms bearing SBS-MM1 containing significantly more mutations than those without the signature (Wilcoxon test, p<0.001, **Extended Data Figure 6a)**. The absence of SBS-MM1 in these tMN can either be explained by malignant progression driven by multiple clones, in parallel (**Extended Data Figure 5b**), or by a cell that managed to escape exposure, outright (**Extended Data Fig 5c**). A lack of effect of disease latency on platinum-associated signature penetrance, and a similar latency between chemotherapy signature-positive and -negative melphalan cases (Wilcoxon test, p = 0.7) suggest the latter model is most likely.

Three lines of evidence further support the model of melphalan escape and reinfusion: firstly, SBS-MM1 was not present in any of the five B-ALL that developed following melphalan/ASCT (**Fig 1d**). Secondly, patients with sequential exposure to platinum and then melphalan/ASCT (2 tMN and 1 B-ALL) had tumors bearing only platinum-related mutational signatures, indicating that a single cell exposed to platinum, but not to melphalan, expanded to clonal dominance; explainable only by escape via the leukapheresis product (**Extended Data Fig 5d-e**). Thirdly, interrogation of the WGS of tumors with precursors that would not have a possible route of escape from melphalan did indeed have SBS-MM1 signatures: a patient with tMN following treatment with oral melphalan without ASCT (IID_H201267) and a transitional cell carcinoma diagnosed following melphalan/ASCT (**Methods, Fig 1d, Extended Data Figure 7**). The latter tumor was chosen for sequencing because unchanged melphalan is partially excreted in urine^26^, thus putting urothelium in direct contact with the mutagen and here providing evidence of clonal expansion of a single melphalan-exposed urothelial cell 8 years following exposure. Our observations suggest that the transplant procedure can allow for a pre-leukemic clone to escape exposure to chemotherapy via leukapheresis and then be reinfused into the patient where it subsequently expands into tMN.

## The genomic driver landscape of therapy-related myeloid neoplasms

Though SNV in driver mutations for tMN and *de novo* AML have been extensively reported^22, 27, 28^, the impact of chemotherapy on driver SNV is still undefined. We first determined positively-selected driver genes using the dN/dS algorithm for both *de novo* AML and tMN, importing samples from the Beat AML dataset to increase power (298 *de novo* AML and 22 tMN; **Methods, Extended Data Fig 8a, Extended Data Table 10**)^28^. The only driver gene with a higher incidence in tMN as compared with *de novo* AML was *TP53* (Fisher test, p = 0.004; FDR = 0.15)^1^. There were no significant differences in mutated driver genes between chemotherapy signature-positive and - negative tMN (**Extended Data Table 11**). However, when the mutated driver genes from all tMN cases with melphalan and platinum exposure were pooled, mutational signature analysis revealed evidence of chemotherapy-associated SBS signature contribution (**Extended Data Figure 8b**). This finding indicates that melphalan and platinum can potentially induce mutations in driver genes of exposed cells.

We next compared recurrent copy number aberrations between *de novo* AML and tMN using GISTIC 2.0 (**Methods**). We differentially detected seven arm-level and one focal region (19p13.2) of amplification and seven arm-level and eleven focal regions of copy number loss (FDR<0.1, **Fig 2a, Extended Data Table 12**). Once tMN samples were stratified by the presence of chemotherapy-induced mutational signatures, a striking pattern emerged: *de novo* AML and tMN lacking chemotherapy-induced mutagenesis (i.e., chemotherapy SBS signature-positive) shared a similar distribution and frequency of copy number events, while tMN harboring chemotherapy signatures were responsible for the majority of significant copy number aberrations, including deletions of chromosomes 5q, 7q, 17p, and focal gain in 19p13.2 (FDR<0.1, **Fig 2a, Extended Data Fig 9a, Extended Data Table 12**).

**Fig. 2.**
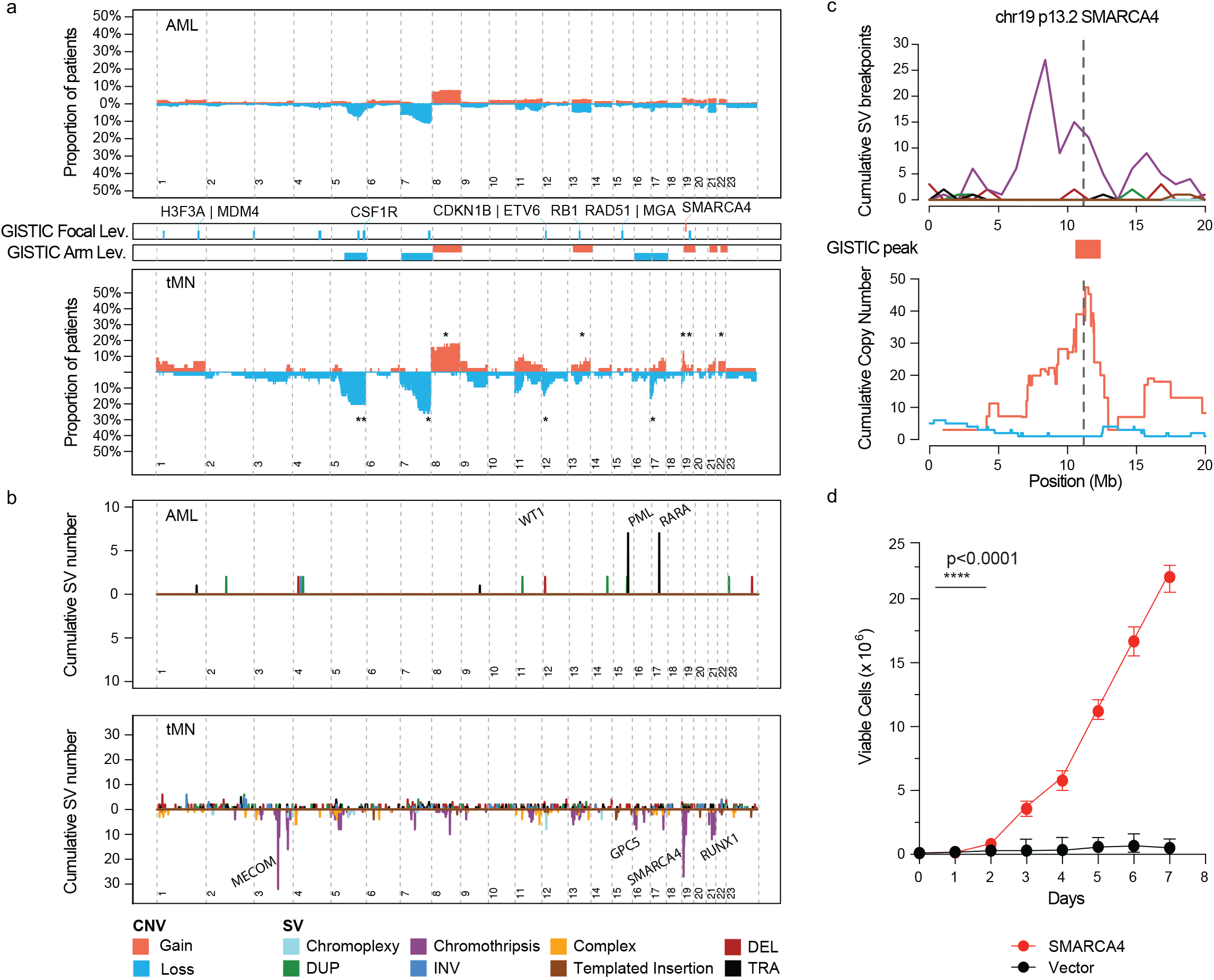
Copy number and structural variant landscapes in therapy-related myeloid neoplasms. **a)** Top. Cumulative copy number profile for all *de novo* AML samples (n = 316; 18 genomes from TCGA and 298 exomes from Beat AML). Bottom. Cumulative copy number for 39 tMN genomes and 22 tMN exomes. GISTIC peaks and CNV arm-level events enriched in tMN (p<0.05, FDR<0.1; Fisher test) are annotated with an asterisk and reported between the two plots. **b)** Structural variant landscape across all *de novo* AML (Top; n=18) and tMN genomes (Bottom; n=39). SV breakpoints are binned into 1 megabase segments. For visual purposes, simple events point upward from the x-axis and complex events (e.g. chromothripsis) point downward. **c)** Plot of cumulative copy number changes and SV chromothriptic breakpoints involving *SMARCA4*. **d)** Growth curves for *SMARCA4* transfected Ba/F3 cells vs. cells transfected with vector in IL3 cytokine independence assay.

To identify the structural mechanisms for copy number aberrations, the landscape of structural variants was interrogated for both *de novo* AML and tMN WGS (**Methods, Fig 2b**). Consistent with the copy number aberrations, complex structural variants (i.e., templated insertions, chromoplexy, and chromothripsis) were enriched in tMN evolving under the influence of chemotherapy-mutagenesis vs those without (Wilcoxon test, p<0.001; **Extended Data Fig 9b**,**c, Extended Data Table 13**). Chromothripsis, a catastrophic shattering and haphazard repair of multiple chromosomal regions resulting in the simultaneous introduction of disparate drivers, was seen in eight of 39 tMN (20.5%)^29-32^. Strikingly, chromothripsis involving chromosome 19p13.2 with focal and multiple amplifications (median 7; range, 5-12 copies) of the *SMARCA4* locus comprised five of these cases with 4 of 5 events found in chemotherapy signature-positive genomes (**Fig 2c, Extended Data Fig 10**). In comparison, across the entire cohort of de novo AML genomes and exomes (n=316), focal amplification of *SMARCA4* was seen in just one case with multiple chromosomal aneuploidies (**Extended Data Fig 11a**). Deleterious mutations in *SMARCA4*, generally considered a tumor suppressor gene, have been implicated in other malignancies including ovarian carcinoma, and various lymphomas^33-35^. As we found it to be uniformly amplified in tMN as a result of complex structural variations, we sought to verify that *SMARCA4* overexpression could promote growth in leukemic cells. *SMARCA4* transfection into Ba/F3 cells indeed demonstrated that gain-of-function amplification could drive growth when compared to vector with IL3 cytokine independence (**Fig 2d, Methods, Extended Data Fig 11b**). This novel dysregulation of the SWI/SNF chromatin-remodeling complex appears a key driver in tMN harboring evidence of chemotherapy-induced mutagenesis. Consistent with previous evidence the *MECOM* locus was also frequently involved by tMN structural variation; with a preponderance for tMN containing a chemotherapy-associated signature (3 of 4 cases, **Fig 2c, Extended Data Fig 10**)^36^.

Pooling together all somatic events among tMN genomes (i.e., structural variants, copy number loss, and SNV), chemotherapy signature-positive cases had a significantly higher prevalence of *TP53* loss (10/16, 62.5%) compared to chemotherapy signature-negative cases (3/23, 13%; Fisher test, p=0.002). Strikingly, among those receiving melphalan/ASCT, all six cases with the SBS-MM1 signature had an event involving *TP53* as compared to 2/10 (20%) without the signature (Fisher test, p = 0.007; **Extended Data Table 14**). Acknowledging the limited sample size, this stark difference indicates that loss of *TP53* may influence the route of expansion following transplant allowing precursors to survive direct exposure to melphalan and accrue chemotherapy-mediated drivers, while precursors with unperturbed *TP53* may preferentially be reinfused in the transplant.

## Chemotherapy-related mutational signatures as molecular barcodes

Several groups have demonstrated that the accumulation of mutations attributable to constant, clock-like processes (SBS1, SBS5, and SBS-HSC) can be used to estimate the age at which a large chromosomal gain occurs in an individual^8, 37-40^. However, unlike in other malignancies, in tMN there is a lack of correlation between clock-like SBS signatures and the age at diagnosis (**Extended Data Fig 12a**)^12^. We therefore leveraged chemotherapy-associated signatures as temporal barcodes by which to time chromosomal gains relative to chemotherapy exposure. Specifically, if a clonal mutation is duplicated across a chromosomal gain, it must have been present prior to the event (i.e., pre-gain). Consequently, a chemotherapy-associated mutational signature present within duplicated mutations necessitates that the exposure occurred prior to the gain. Conversely, chemotherapy-associated mutational signatures present only among nonduplicated clonal or subclonal mutations implies that the exposure occurred after the chromosomal gain (i.e., post-gain, **Fig 3a**)^41^.

**Fig. 3.**
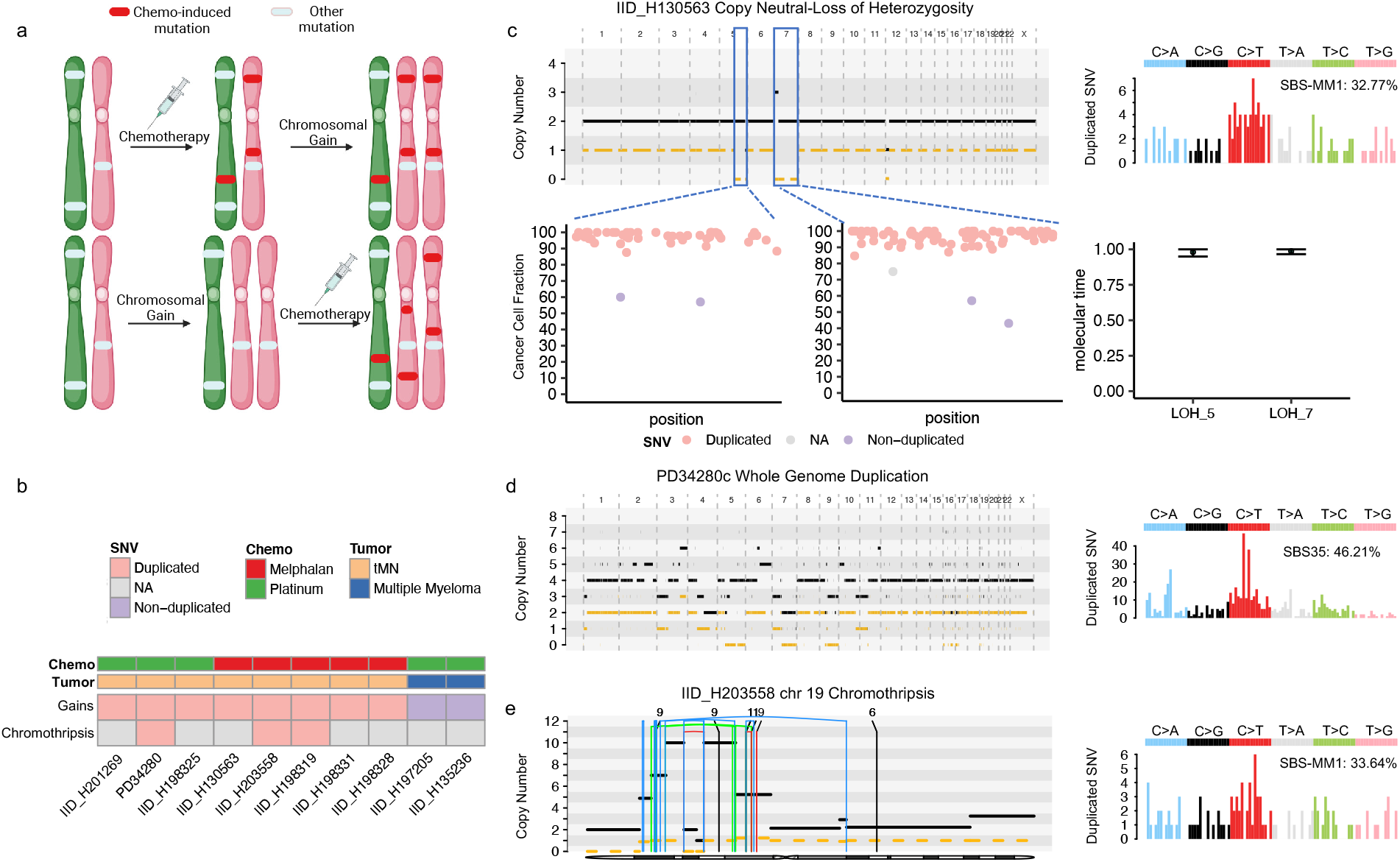
Chemotherapy-associated single base substitution signatures as temporal barcodes. **a)** Cartoon summarizing the rationale used to time chromosomal duplications according to the chemotherapy exposure. **b)** Heatmap revealing the pattern of timing for chromosomal duplications and chromothripsis events with relation to chemotherapy for 8 tMN and for two cases of post-chemotherapy multiple myeloma. **c)** Example of two copy neutral loss of heterozygosity (CN-LOH) acquired after the chemotherapy exposure. In this case, both CN-LOH contain duplicated mutations (bottom left) consistent with late acquisition as confirmed by molecular time analysis (bottom right) and showing large SBS-MM1 (melphalan) signature contribution within duplicated (i.e., pre-gain) mutations (top right). **d)** Example of whole genome duplication in a tMN relapse (left). Duplicated mutations (right) showed a large SBS35 (platinum) contribution, suggesting this event was acquired after the platinum exposure. **e)** Chromothripsis event on chromosome 19 (*SMARCA4*) with multiple duplications (left). Similarly to (**d**), the duplicated mutations mutational signature contribution within chromothripsis-associated amplifications (right) were enriched for SBS-MM1 contribution, implying that the chromothripsis event was acquired after the melphalan exposure. In (**c-e**) The horizontal black line indicates the total copy number; the dashed orange line indicates the minor copy number. In (**e**) the vertical lines represent SV breakpoints, color-coded based on SV class: blue = inversion, green = tandem-duplication; red = deletion; black = translocation.

After collapsing together large events (i.e., trisomy, copy neutral loss of heterozygosity, and whole genome duplication) that occurred within the same time window (**Methods, Extended Data Fig 12b**), eight tMN with chemotherapy-associated SBS signatures were amenable to chemotherapy barcoding. Strikingly, in all cases, the melphalan or platinum signatures were detectable within duplicated clonal mutations implying that large copy number aberrations occurred during or after exposure to chemotherapy and late in tMN evolution (**Fig 3b**,**c**,**d, Extended Data Table 15**). As all copy number aberrations associated with chromothripsis occur simultaneously within one catastrophic event^42^, we similarly applied this methodology to three tMN with large chromothriptic events characterized by amplified genomic segments with more than 40 clonal non-clustered mutations^8, 40^. The chemotherapy-associated signatures were present in the duplicated mutations, including the previously mentioned events on *SMARCA4*, supporting that these complex structural variants occurred after exposure to mutagenic therapy (**Methods, Fig 3b, e, Extended Data Table 16**).

In contrast to tMN, for two multiple myeloma cases occurring post-platinum exposure for an unrelated primary solid tumor, the associated platinum signatures were seen only in subclonal (i.e, non-duplicated) mutations, in agreement with prior knowledge that many chromosomal gains are initiating and early events (**Fig 3b**) ^8^. We sought to incorporate the known absolute timing of chemotherapy exposure into the context of the cancer-initiating events for these two multiple myeloma patients to further validate our approach of chemotherapy-associated SBS signatures for molecular timing. We first calculated the individual SBS5 mutation rate per year for the two secondary tumors and found no significant difference when compared to 77 WGS from 47 patients with primary or smoldering multiple myeloma (**Methods, Extended Data Fig 12c**)^8^. Collapsing together large chromosomal gains occurring within the same time window, we estimated the SBS5-based molecular time in order to predict at which age these two patients acquired the first multi-chromosomal gain event and the emergence of the most recent common ancestor (**Methods, Fig 4a, Extended Data Figure 12b**). Consistent with prior reports, in both cases, the initiating multi-chromosomal gain events were estimated to have occurred in the 2nd decade of life^8^ and the most recent common ancestor was timed to having arisen prior to diagnosis of the unrelated primary malignancy (i.e., the solid tumor) and the associated platinum exposure. This SBS5-based molecular timing was in agreement with the presence of platinum-associated chemotherapy signatures in subclonal mutations. Overall, chemotherapy-associated mutational signatures are thus temporal molecular barcodes that can be used to time the acquisition of large-scale copy number duplications over a patient’s life.

**Fig. 4.**
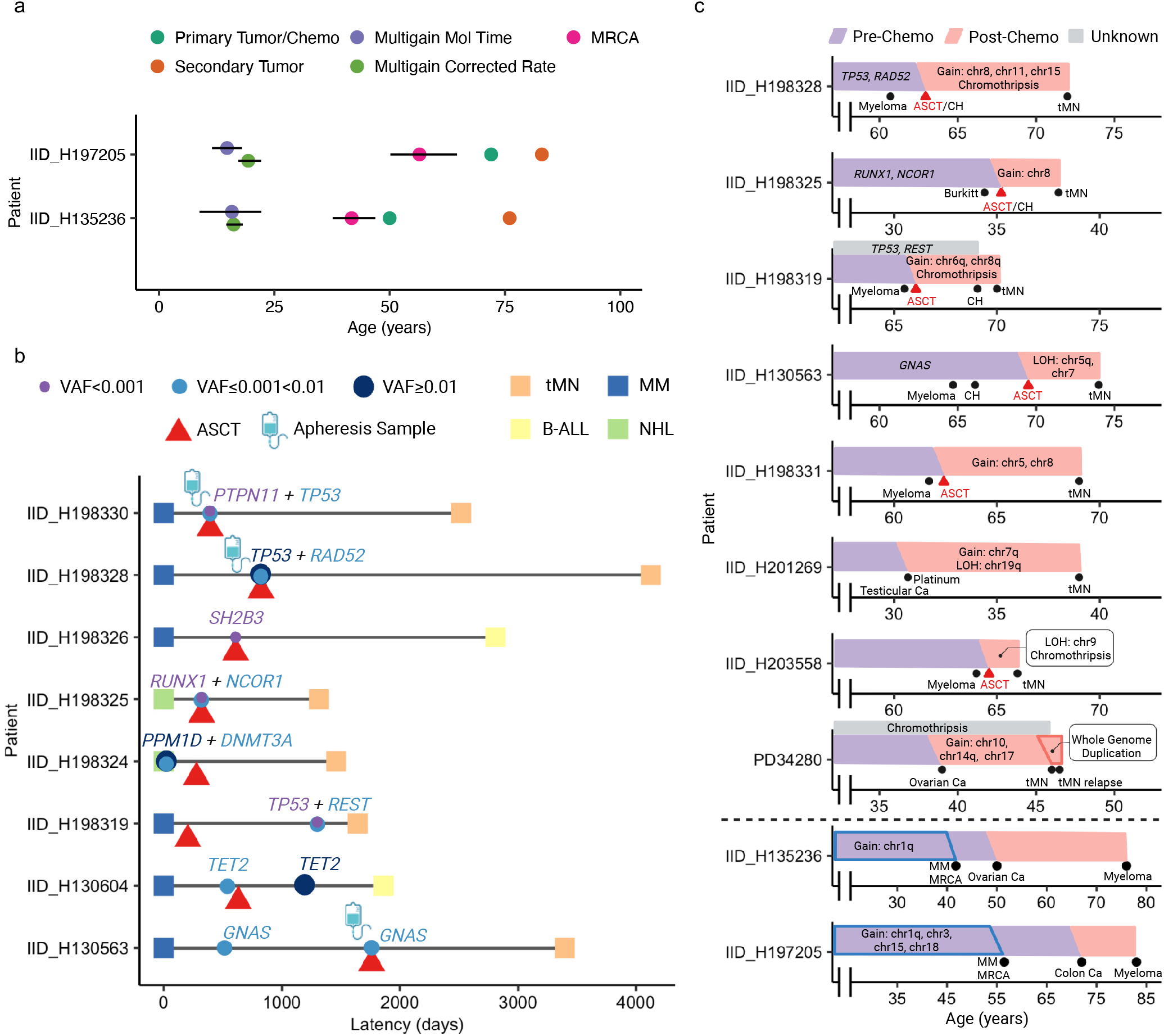
Evolutionary history of chemotherapy-exposed malignancies. a) Absolute timing estimates of large chromosomal gain acquisition in two platinum-exposed multiple myeloma with 95% Confidence Intervals. Multi-gain events for multiple myeloma were timed with two orthogonal techniques (**Methods**): one using the SBS5-based molecular time (i.e., multi-gain mol time), and the other using the individual SBS5 mutational rate per year estimated by a linear mixed effect model (i.e., multi-gain corrected rate). Platinum was administered directly following primary tumor diagnosis. MRCA: most recent common ancestor. **b)** Swimmer plot depicting timing of clonal hematopoiesis assessment in relation to malignant diagnoses and transplant dates. Latency reflects time from diagnosis of primary tumor until diagnosis of therapy-related malignancy. All samples were collected prior to melphalan administration aside from IID_H198319. **c)** Reconstruction of tumor evolution for select cases using antecedent clonal hematopoiesis, molecular time, and duplicated chemotherapy mutation data for tMN (top) and multiple myeloma (bottom).

## Detection of Antecedent Clonal Hematopoiesis

To further reconstruct the genomic life-history of tMN, we next performed target capture-based sequencing with a bait panel of 505 cancer genes (MSK-IMPACT) to a median depth of 755x on pre-melphalan blood mononuclear cells, granulocytes, or CD34+ apheresis samples from 11 hematologic malignancies, all of which were from patients with multiple myeloma treated with melphalan/ASCT (**Methods**)^17, 43^. For eight of the eleven cases (72%), including three of four leukapheresis products, evidence of antecedent pre-leukemic clones was detected via somatic variants that subsequently became clonal in tMN (**Fig4b, Extended Data Table 17**). Notably, for IID_H198330, a case without the melphalan mutational signature, an antecedent *TP53*-mutated CH was detected in the apheresis product (**Extended Data Fig 13**), supporting an evolutionary trajectory of escape from exposure to chemotherapy, reinfusion with transplant, and expansion in the absence of melphalan-induced mutagenesis.

## Discussion

Using WGS of tMN samples from patients exposed to a variety of anti-neoplastic agents and targeted sequencing of pre-therapy samples, we characterized the mutational impact of chemotherapy, and used chemotherapy-induced SBS signatures as a temporal barcode to reconstruct the evolution of tMN from its antecedent clonal hematopoiesis (**Fig 4c**). While tMN evolving without chemotherapy-induced mutagenesis have features like those of *de novo* AML, tMN with chemotherapy-induced mutagenesis (i.e., chemotherapy mutational signatures) are relatively hypermutated and more likely to harbor complex genomes inclusive of chromothripsis and copy number aberration. These data suggest that exposure to mutagenic chemotherapeutics such as platinum and melphalan not only introduce hundreds of somatic mutations, but also promote genomic complexity and the selection of distinct genomic drivers. This investigation also highlights that WGS is needed to characterize the full landscape of alterations in these complex malignancies.

Due to the lack of correlation between clock-like mutational processes and age, it has previously not been possible to estimate the absolute time at which certain driver events were acquired in tMN development^12^. To overcome this historical limitation, we leveraged chemotherapy-associated mutational signatures as temporal barcodes linked to discrete clinical exposures to demonstrate that complex events occur late in tMN evolution, following chemotherapy exposure. This approach also allowed to perform an orthogonal validation of previously reported molecular timing approaches for determine the age of large chromosomal gains^8, 38^.

Finally, we demonstrated that patients with bona fide CH who undergo ASCT can develop tMN that bear no evidence of melphalan-induced mutations (i.e., SBS-MM1 mutational signature). Although previously known that an increased chemotherapy-induced mutational burden is not requisite for tMN progression^12, 18^, post-ASCT tMN expansion in the absence of a melphalan signature indicates that not even direct cellular exposure to chemotherapy is needed. This route to malignant transformation places increased emphasis on potential leukemia-permissive effects of chemotherapy on the bone marrow compartment^17, 18, 44^. Furthermore, it appears that *TP53* status may influence the trajectory of post-ASCT tMN development with regard to whether precursors may survive direct exposure to high-dose melphalan and its resultant mutagenesis or preferentially be reinfused to circumvent exposure.

In compatibility with our findings, chemotherapy and transplantation have been shown to confer selective pressure on the remaining progenitor cells in a relatively vacant marrow niche, such that hematopoiesis is reconstituted by a limited number of clones^45-48^. In addition, high rates of secondary myeloid neoplasms have been observed following novel chimeric antigen receptor T-cell therapy (CAR-T), which is also known to be associated with prolonged immune suppression^49-51^. Altogether, several lines of evidence suggest that the evolution of precursors states into their malignant successors is likely driven by a complex interaction between inflammation, mutagenesis, and immune suppression^14, 18, 52, 53^. As more immunomodulatory and cellular therapies move into widespread clinical practice, and as patients live longer following therapeutic advancements, it will be increasingly important to characterize how this complex interplay is affected by these therapies to promote tMN evolution.

## METHODS

### Study cohort

The WGS cohort was compiled with newly sequenced and publicly available data (**Extended Data Table 1**). Clinical records at Memorial Sloan Kettering Cancer Center were screened to identify adult patients who had developed tMN following exposure to either of melphalan-or platinum-containing anti-neoplastic regimens. 18 tMN were eligible for sequencing (**Extended Data Tables 1-2**). The remaining 22 tMN genomes (from 21 patients) were imported from public datasets^18, 23^ (dbGaP: phs000159 and EGAD00001005028). Additionally, non-myeloid secondary malignancies were sequenced for the study: clinical records at Memorial Sloan Kettering Cancer Center were screened to identify patients with hematologic and solid tumors that developed following exposure to melphalan and platinum. Specifically, 5 patients with B-ALL following melphalan/ASCT, one patient with a secondary bladder tumor following exposure to melphalan/ASCT and two patients with secondary multiple myeloma following treatment with platinum-based chemotherapy had secondary tumors eligible and selected for WGS. All samples and data were obtained and managed in accordance with the Declaration of Helsinki and the Institutional Review Board of Memorial Sloan Kettering Cancer Center. The study involved the use of human samples, which had been collected after written informed consent had been obtained. 21 *de novo* AML whole genomes (including three relapse samples) were imported from a TCGA dataset^22^(dbGaP 000178) as comparators to tMN. 298 *de novo* AML and 22 tMN whole exomes were imported from the Beat AML dataset (dbGaP: phs001657).

### Sample preparation

Tumor samples for in-house sequencing were curated from three sources. Tumors from high purity samples (and solid tumors) that had previously had clinical molecular testing performed and had leftover cDNA directly sequenced. The two secondary multiple myeloma samples were sequenced directly from leftover DNA extracted from CD138-sorted plasma cell used for clinical SNP-array. For the remaining, frozen bone marrow mononuclear cell aliquots were obtained from an in-house biobank and were either sequenced directly following DNA extraction or were first sorted via fluorescence-activated cell sorting to improve sample purity. Matched normal were selected from frozen peripheral blood mononuclear cells, peripheral blood granulocytes, or CD34-selected autograft products were used as normal match (**Extended Data Table 1**).

### Whole-genome sequencing

Following quantification via PicoGreen and quality control by Agilent Bioanalyzer, ∼500 ng of genomic DNA was sheared (LE220-plus Focused-ultrasonicator; Covaris, catalog no., 500569) and sequencing libraries were prepared using a modified KAPA Hyper Prep Kit (Kapa Biosystems, KK8504). Briefly, libraries were subjected to a 0.5 × size select using aMPure XP Beads (Beckman Coulter, catalog no., A63882) after post-ligation cleanup. Libraries that were not amplified by PCR (07652_C) were pooled equivolume. Libraries amplified with five cycles of PCR (07652_D, 07652_F, and 07652_G) were pooled equimolar. Samples were run on a NovaSeq 6000 in a 150 bp/150 bp paired-end run, using the NovaSeq 6000 SBS v1 kit and an S4 Flow Cell (Illumina), as described previously^54^. Target coverage depth was 70x for tumor and 40x for normal.

### Whole-genome analysis pipeline

Coverage for tumor and normal samples are reported in **Extended Data Table 1**. Short insert paired-end reads were aligned to the reference genome (GRCh37) using the Burrows–Wheeler Aligner (v0.5.9; ref. 17). All samples were uniformly analyzed by the following bioinformatic tools: somatic mutations were identified by CaVEman^54^; copy number analysis and tumor purity (i.e., cancer cell fraction) were evaluated using Battenberg (https://github.com/Wedge-Oxford/battenberg); SVs were defined by BRASS (https://github.com/cancerit/BRASS) via discordant mapping of paired-end reads, passed through additional quality filters, and were manually curated to define complex events (i.e., templated insertions, chromothripsis, and chromoplexy) as described previously. The phylogenetic tree of each case was reconstructed using Pyclone-VI (https://github.com/Roth-Lab/pyclone-vi) to determine clonal and subclonal variants.

The exomes data downloaded from the public repository were aligned to the reference human genome (GRCh37) using Burrows-Wheeler Aligner, BWA (v0.7.17). Deduplicated aligned BAM files were analyzed using FACETS (v0.5.6, https://github.com/mskcc/facets) for copy number variants, CaVEMan (v1.13.14, https://github.com/cancerit/dockstore-cgpwxs) for single nucleotide variants (SNVs) and Pindel (v3.2.0, https://github.com/cancerit/dockstore-cgpwxs) for small insertions-deletions.

The genome regions that were significantly modified in our cohort were identified by using GISTIC2.0 (v2.0.23, https://www.genepattern.org). To improve the test’s statistical power, we ran our cohort of myeloid whole genomes (n=57; not including relapse cases) with Beat-AML samples (n=320). In this way we were able to detect the anomalous peaks and arms shared among all the sample. The analysis was executed using Gene Pattern web interface (http://genepattern.broadinstitute.org) and setting a q value threshold of 0.01. For further comparison, samples were split by status as *de novo AML*, tMN with chemotherapy mutational signature (i.e., chemotherapy-induced mutagenesis), and tMN without chemotherapy mutational signatures. Because mutational signatures were not run for exomes, Beat-AML tMN samples were excluded from this final comparison (n=22).

We applied the dN/dScv method to detect genes under positive selection in our cohort^55^. To increase the statistical power, we included 320 *de novo* AML and tMN samples from the Beat-AML study.

### Mutational Signatures

Mutational signatures were analyzed across all whole genomes. To estimate the activity of mutational signatures, we first employed a three step process of de novo extraction, assignment, and fitting^23^. For the first step, we ran SigProfiler for SBS, DBS, and ID signatures^24^. All extracted signatures were then compared with the latest Catalogue of Somatic Mutations in Cancer (COSMIC) reference (https://cancer.sanger.ac.uk/cosmic/signatures/SBS) to identify the known mutational processes active in the cohort. In the case of ID signatures, the deconvolution and fitting solution was accepted outright. For SBS and DBS signatures, we required the addition of signatures not currently included in the most recent version (3.2) of the COSMIC catalogue (**Extended Data Table 4**). These are SBS-MM1 (melphalan)^8^, SBS-HSC (clock-like signature in hematopoietic cells)^12^, E-DBS3 and E-DBS9 (platinum)^6^. We performed an adjusted deconvolution with the respective SBS and DBS COSMIC catalogues with the addition of these four signatures using a bespoke algorithm (https://github.com/UM-Myeloma-Genomics/Signature-Assignment)^23^. The code generates a pairwise fitting contribution of user-supplied reference mutational signatures to *de novo* extracted signatures and is particularly useful for the addition and evaluation of signatures not included in the COSMIC reference. The top deconvolution combination with biologic rationale reflective of signatures known to be active in included tumor histologies was chosen for each *de novo* signature extraction unless the SigProfiler solution was more appropriate. Deconvolution revisions are marked with an asterisk in **Extended Data Figures 2-3** and reported in **Extended Data Tables 6 and 8**^23^. For SBS, we applied mmsig (https://github.com/UM-Myeloma-Genomics/mmsig)^56^, a fitting algorithm, to confirm the presence and estimate the contribution of each mutational signature in each sample guided by the catalog of signatures extracted for each individual sample by SigProfiler, our revised deconvolution, and with revisions for processes known or not known to be active in disease histologies (i.e., addition of SBS8 for multiple myeloma samples, removal of flat SBS40 signature in all cases in favor of SBS5 and SBS-HSC). mmsig confidence intervals were generated by bootstrapping 1,000 mutational profiles from the multinomial distribution each time repeating the signature fitting procedure, and finally taking the 2.5th and 97.5th percentile for each signature. At least 40 mutations were required per sample for this analysis.

### Chemotherapy-Related Mutational Signatures in Transitional Cell Carcinoma and in tMN Treated with Oral Melphalan

SBS signature de novo extraction for a single sample is technically ill-advised. We imported mutational signatures for the PCAWG cohort of Transitional Cell Carcinoma (n=23)^24^ and then we used MutationalPatterns plot_compare_profiles and cos_sim functions (https://github.com/UMCUGenetics/MutationalPatterns) to compare the case 96-profile with de novo signatures extracted by Sigprofiler. The difference in mutations was quantified and then compared directly to the SBS-MM1 mutational signature using cosine similarity (**Extended Data Figure 7b**). A similar approach was applied for tMN exposed only to oral melphalan in the absence of ASCT (**Extended Data Figure 7a**), in which SBS-MM1 was not previously detected^12,18^. Mutational signatures were first extracted in the tumor and then for 18 *de novo* AML in the cohort. The difference in mutational profiles was then ascertained and compared to SBS-MM1 using cosine similarity.

### Cell culture, *SMARCA4* transfection and cytokine independence assay

Ba/F3 cells were gifted to Dr. Taylor from Dr. Omar Abdel-Wahab (Memorial Sloan Kettering Cancer Center). The cells were cultured in RPMI media supplemented with 10% FBS, 1% Penicillin/Streptomycin and 10 µg/mL of mouse IL3 (mIL3) and were maintained at 37°C and 5% CO_2_. pQCXIH *BRG1* was a gift from Joan Massague (Addgene plasmid # 19148; RRID:Addgene_19148). The *BRG1/SMARCA4* plasmid was linearized with SalI and transfected into Ba/F3 cells via electroporation (Neon transfection system, ThermoFisher, Waltham, MA). Hygromycin was utilized to select for *SMARCA4* overexpressing Ba/F3 cells and overexpression was confirmed by immunoblotting with BRGI/SMARCA4 antibody (Cell Signaling E906E, 1:500 dilution). *SMARCA4* or vector expressing Ba/F3 cells were then cultured in media without IL3. Cells were seeded in triplicates at a starting concentration of 100,000 cells/mL and were counted daily using Vi-Cell BLU automated cell counter (Beckman Coulter, Indianapolis, IN) and plotted using GraphPad Prism Version 9 Software.

### Molecular Time and Absolute Timing of Gains

The relative timing of large chromosomal duplications (multi-gain events) was estimated with the mol_time function (https://github.com/UM-Myeloma-Genomics/mol_time). As previously described, this approach allows for the relative timing of gains of large chromosomal segments^8, 38^ by using the corrected ratio of clonal single nucleotide variants duplicated or non-duplicated across the gains: VAF 66% if duplicated and found on two alleles (pre-gain mutation) and VAF 33% if non-duplicated and found on one allele (post-gain mutation). VAFs were corrected for sample purity (i.e., cancer cell fraction) by combining Battenberg’s estimation of tumor purity and the density and distribution of SNV VAF within clonal diploid regions of each sample genome. We required that gains have a minimum of 40 clonal mutations to be included in analysis for accuracy concerns^8^. For the purposes of increasing power to accurately detect mutational signatures, gains occurring in an overlapping time window estimate were collapsed (i.e, multi-gain event within the same molecular time). For myeloid cases, this provided evidence of relative timing of gain with respect to tumor diagnosis and corroborated results of the following duplicated mutation analysis.

For two multiple myeloma cases, mutational signatures were quantified for pre-and post-gain mutations. The mutational burden of clock-like mutations (SBS5) was then used to ascertain each patient’s individual SBS5 mutation rate. This was accomplished by pooling the two secondary multiple myeloma tumors and the multiple myeloma longitudinal cohort from Rustad et al., 2020 to use a linear mixed effects model to estimate SBS5 accumulation over time^8^. Then, to convert molecular time estimates into absolute time: i) the most recent common ancestor was estimated dividing clonal SBS5 mutations by the individual SBS5 mutation rate with interval of confidence derived from the upper and lower bounds of standard deviation for the patient-specific mutation rate; ii) the multi-gain event was estimated by dividing the number of clonal SBS5 pre-gain mutations by the mutation rate corrected for size of the gains and the interval of confidence derived from the mutation rate as above; iii) the multi-gain event was additionally estimated by multiplying the MRCA by the molecular time estimate with interval of confidence generated from bootstrapping the molecular time estimate 1000 times. Only gains with at least 40 clonal mutations were included for the purpose of accuracy.

### Duplicated Mutational Burden and Chemotherapy Barcoding

In an approach similar to that used for estimated timing of gains using duplicated clock-like mutations^8, 38^, we leveraged mutations that were duplicated across gains (i.e., VAF 66%) to determine variants that predated the copy number gain. VAFs were corrected for sample purity (i.e., cancer cell fraction) by combining Battenberg’s estimation of tumor purity and the density and distribution of SNV VAF within clonal diploid regions of each sample genome. Mutations from within different gains were collapsed if molecular time (above) supported the gains as occurring in the same time window. For IID_H198325, IID_198331, and IID_H198328 the molecular time was not possible to be estimated because of either low mutational burden or excess of unassigned mutations. Given the patterns observed in the other tMN, the gains from these samples were treated as having occurred in the same molecular time for the purposes of this approach.

Tumor phylogeny from Pyclone, above, revealed clonal and subclonal mutations and all SNV were subsequently grouped into clonal duplicated and non-duplicated groups, and subclonal (non-duplicated) groups. Mutational signature analysis was run on each group of intra-gain mutations, as above, to determine pre-and post-gain mutational processes active before and after the gain. At least 40 total mutations per multi-gain were required in a group to confidently ascertain mutational signature contribution.

A similar approach was applied to amplifications that were caused by chromothripsis events^42^. After correction of VAF for sample purity, mutations across gains deemed part of chromothripsis events^30^ were pooled and separated into duplicated and non-duplicated groups. Clustered mutations (i.e., kataegis) were filtered out so as not to skew results towards APOBEC mutational signatures^23^. Mutational signature analysis was performed for events with 40 or more clonal mutations.

Clinical correlation with timing of chemotherapy and associated signatures was then compared to molecular time estimates and according to duplication status to reconstruct evolutionary timelines relative to chemotherapy exposure.

### Target capture-based sequencing

Peripheral blood samples and/or CD34-selected autograft products collected prior to chemotherapy exposure were collected from eleven patients with matched whole genomes (**Extended Data Table 17**). Where possible, apheresis product was prioritized. Samples were sequenced using MSK-IMPACT, a Food and Drug Administration-authorized hybridization capture-based next-generation sequencing assay of protein-coding exons from 505 known cancer-associated genes (**Extended Data Table 17**)^17, 43^. Matched normal was obtained by pooling IMPACT-505 data from 8 healthy individuals.

### Target capture-based sequencing variant calling and filtering

Given the potential for extremely low VAF (i.e., small clone size) for CH mutations in normal samples preceding tumor expansion by many years, population-based CH screening techniques^17^ would fail to capture many low allelic frequency variants of interest and so directed mutation query was employed. Our approach consisted of two distinct strategies to characterize antecedent CH variants. Our first approach was targeted-sequencing-centric and was a modified workflow from Bolton et al^17^. First, stringent quality filters were applied to calls from a triple caller pipeline of Mutect, Strelka, and Caveman^54, 57, 58^. We required a SNV to be called by two or more callers, have a VAF of >0.02, have passed default quality flags, and not result in synonymous substitution. We further required at least 10 supporting forward and reverse reads in Mutect, and to further filter any germline SNPs, we removed any variants reported for any population in the gnomAD database at a frequency greater than 0.005. Indels were called with Pindel ^59^ and considered if they passed all quality flags. Further postprocessing filters to remove sequencing artifacts were employed as per Bolton et al.^17^ However, as we also had whole genomes sequencing data, we were also able to work in the reverse direction: for the list of all nonsynonynous mutations and indels identified in driver genes, each individual variant was queried directly in targeted sequencing bam files using Integrated Genome Viewer^60^. If at least one read for the mutation was identified, the mutation was considered present in the target sequencing sample. Variants were further confirmed and VAF was calculated by generating a pileup of reads for the targeted sequencing regions with SAMtools^61^ for both reference and alternate alleles.

## ACKNOWLEDGEMENTS

This work was supported by the International Myeloma Foundation 2020 Brian D. Novis Research Award, by the Paula and Rodger Riney Foundation, by the Sylvester Comprehensive Cancer Center NCI Core Grant (P30 CA 240139) by the Memorial Sloan Kettering Cancer Center NCI Core Grant (P30 CA 008748), by the Conquer Cancer Young Investigator Award (16587), and by the Sylvester Comprehensive Cancer Center/American Cancer Society IRG grant. F.M. is supported by the American Society of Hematology. B.D. is supported by Myeloma Crowd and the Sylvester K12 Calabresi Clinical Oncology Research Career Development Program. N.B. is supported by the Associazione Italiana per la Ricerca sul Cancro (AIRC) thorugh an investigator grant n. 25739

## AUTHOR CONTRIBUTIONS

F.M. designed and supervised the study, collected, and analyzed the data and wrote the paper. O.L. designed the study, collected the data and wrote the paper. B.D. and B.Z. supervised the study, collected, and analyzed the data and wrote the paper. K.M. collected and analyzed the data and wrote the paper. E.P. designed and collected the data. J.T. collected and analyzed the data. E.B. and G.M. collected the data and performed all the flowcytometry sorting experiments. A.Mc., D.Co., and J.A.O. analyzed the data. J.J., M.A., T.M.T performed the SMARCA4 wet validation. S.U., S.N., J.T., M.R., C.H., Y.Z., D.C., A.L., M.S., K.G., H.L., J.H.P, N.B., S.X.L., N.B., J.W. collected the data.

All authors read, revised, and proofed the manuscript

## Data Availability

The dataset used for this paper is derived from public sources: 22 tMN genomes (from 21 patients) were imported from the following public datasets: dbGaP: phs000159 and EGAD00001005028). 21 *de novo* AML whole genomes (including three relapse samples) dbGaP phs000178; 298 *de novo* AML and 22 tMN whole exomes were imported from dbGaP: phs001657.

Data from newly sequenced samples will be made available for public use on dbGaP upon publication of the manuscript.

## Code Availability

Analyses conducted using previously published code are detailed and available via github links in Methods.

