## Supplementary Figures for "Chemotherapy Signatures Map Evolution of Therapy-Related Myeloid Neoplasms"

### Extended Data Figures

**Extended Data Figure 1. Clinical summary of therapy-related neoplasms included in this study. a)** Sankey plot for relationship between primary tumor, therapy, and secondary malignancy whole genome samples. For visual purposes, secondary multiple myeloma and transitional cell carcinoma are not included. **b)** Kaplan-Meier curves for therapy-related AML and MDS from the whole genome cohort.

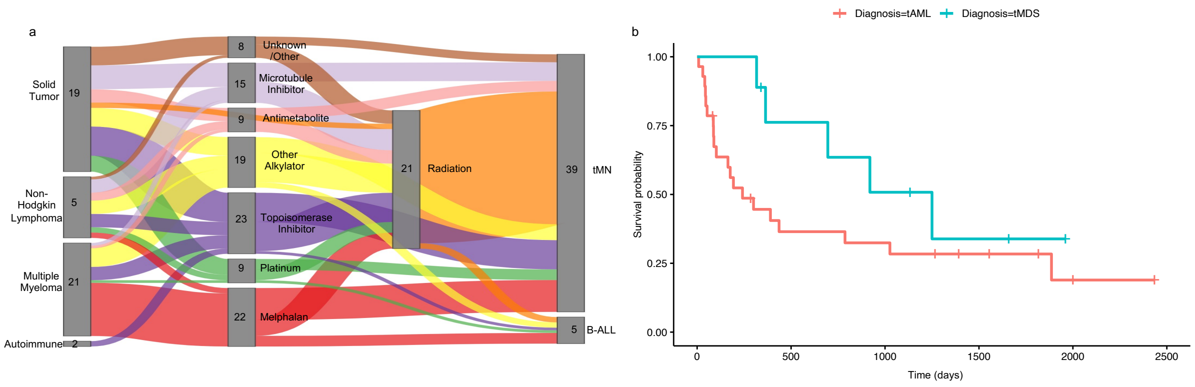

**Extended Data Figure 2. Single Base Substitution (SBS) signatures.** Output from *SigProfiler* SBS signatures de novo extraction. SBS signatures annotated with an asterisk have been revised using a deconvolution solution containing non-COSMIC mutational signatures (**Methods**). The remaining are original *SigProfiler* suggested solutions.

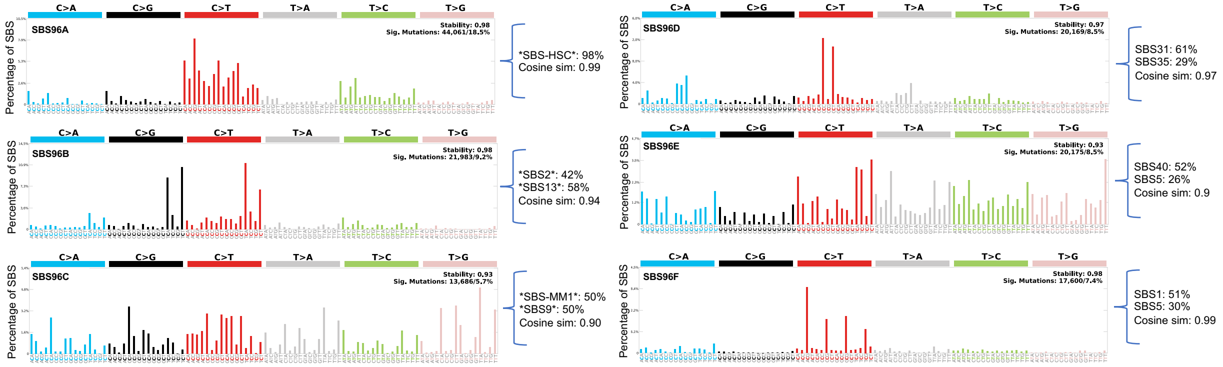

**Extended Data Figure 3. Double Base Substitution (DBS) signatures.** Output from *SigProfiler* DBS signatures annotated with an asterisk have been revised using a deconvolution solution containing non-COSMIC mutational signatures (**Methods**). The remaining are original SigProfiler suggested solutions.

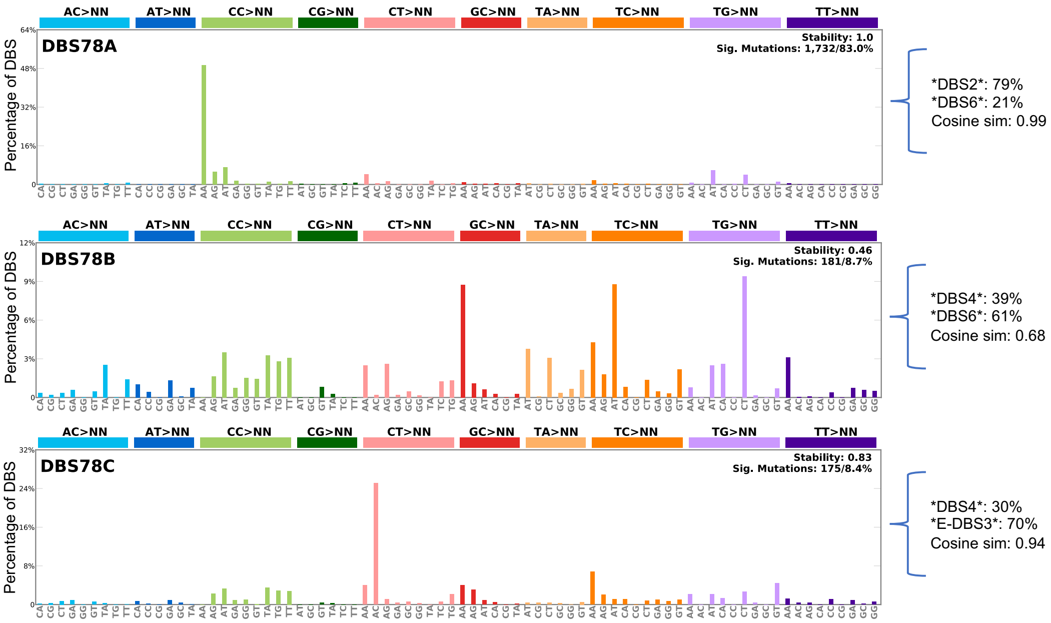

**Extended Data Figure 4. Indel Substitution (ID) signatures.** Output from *SigProfiler* ID signatures extraction with *SigProfiler* suggested deconvolution solutions.

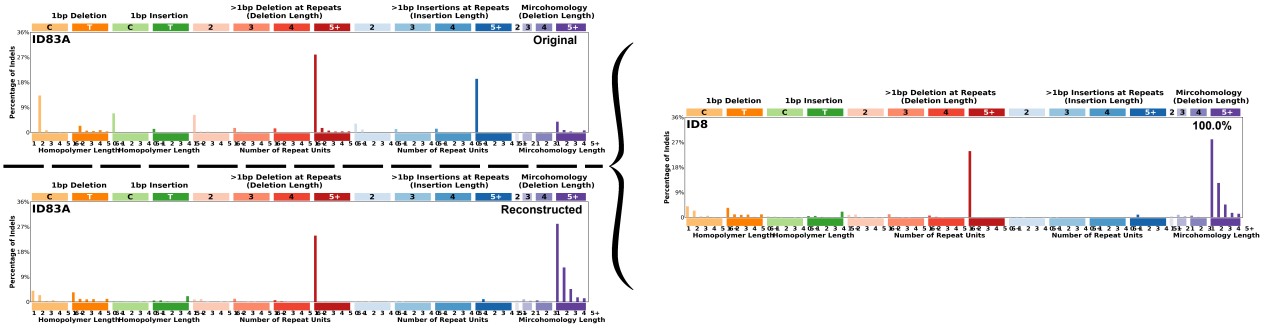


**Cosine Similarity: 0.549** **L1 Error %: 120.0%** **KL Divergence: 2.0751**
**Correlation: 0.51** **L2 Error %: 96.791%**

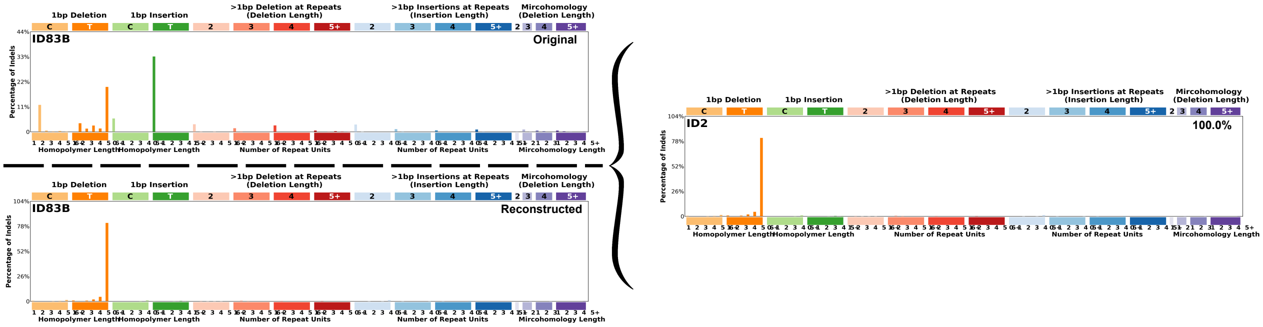


**Cosine Similarity: 0.48** **L1 Error %: 149.0%** **KL Divergence: 7.4872**
**Correlation: 0.465** **L2 Error %: 173.435%**

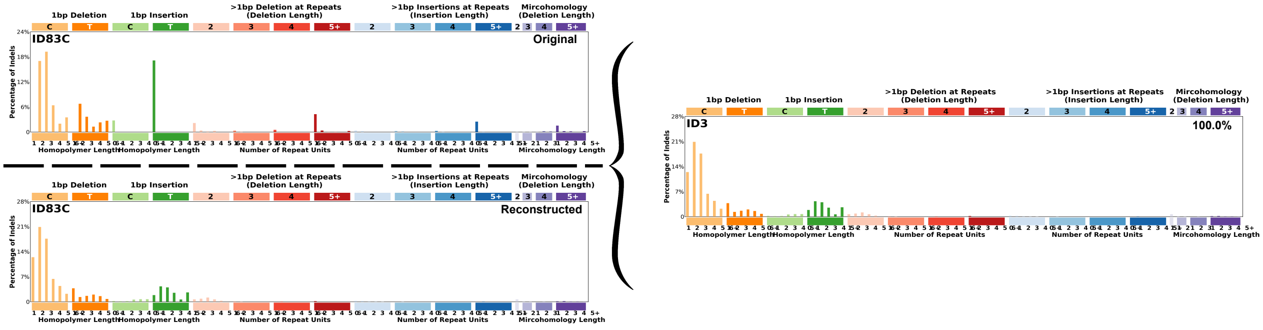


**Cosine Similarity: 0.747** **L1 Error %: 86.0%** **KL Divergence: 1.1266**
**Correlation: 0.715** **L2 Error %: 68.604%**

**Extended Data Figure 5. Schema for the measurement of chemotherapy-related mutational signatures in bulk whole genome sequencing data. a)** Measurement of chemotherapy-associated mutational signatures depends on a single cell, bearing a unique chemotherapy-associated mutational catalogue, to expand to clonal dominance (single cell expansion model). **b)** Polyclonal expansion following chemotherapy exposure does not yield a measurable chemotherapy-associated signature. **c)** Another explanation for lack of signature expression is escape from exposure entirely via leukapheresis and reinfusion. Rather than observing both platinum and melphalan signatures following sequential exposure in the same patient **(d)**, only platinum signatures are present suggesting escape to subsequent melphalan exposure via leukapheresis **(e)**.

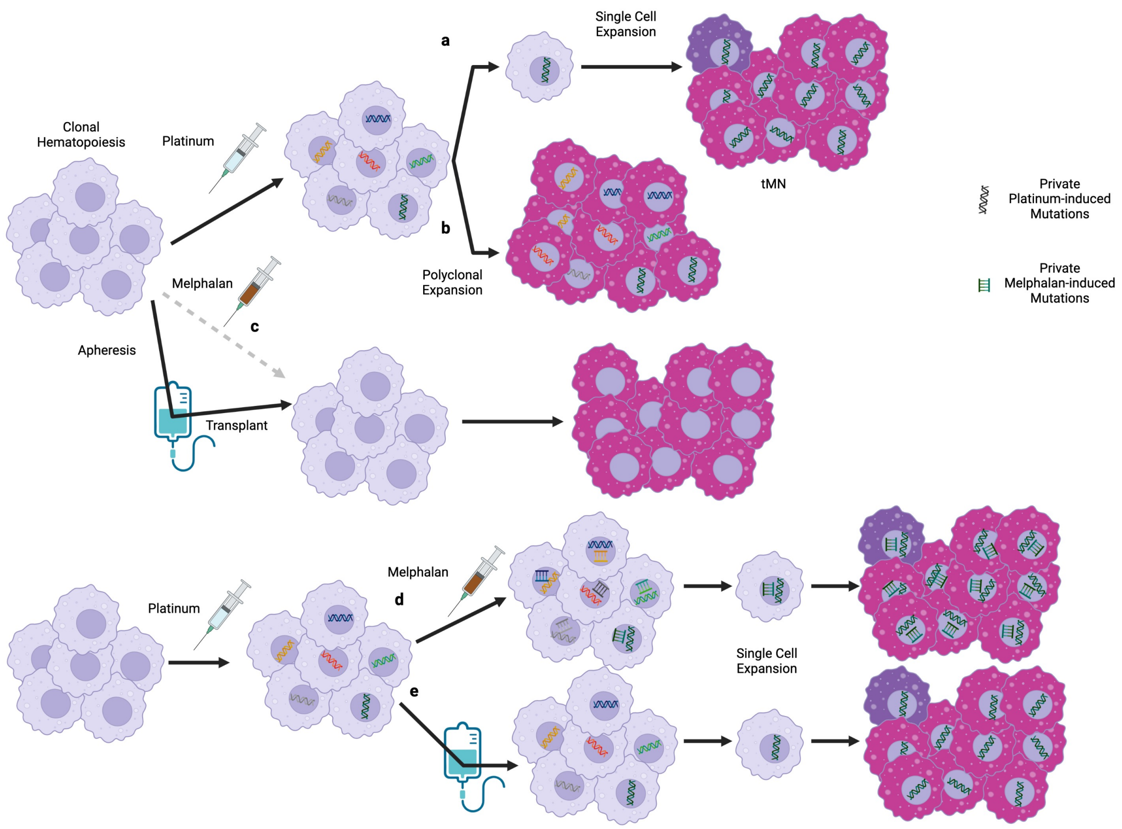

**Extended Data Figure 6. Comparison of SBS mutational burden in post-melphalan therapy-related myeloid neoplasms with or without the SBS-MM1 signature.** The p-value was estimated using Wilcoxon test.

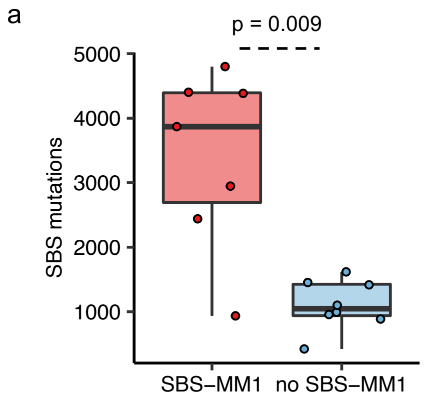

**Extended Data Figure 7. Detecting the SBS-MM1 (melphalan) signature in two tumors without potential for transplant-mediated escape**. **a)** SBS difference between mutational profile for a tMN developing after oral melphalan exposure (IID_H201267) and the cumulative profiles of n=18 *de novo* AML WGS (top). The difference (removing negative contributions) compared to the SBS-MM1 mutational signature profile (bottom). **b)** SBS difference between mutational profile for a melphalan-exposed transitional cell carcinoma (TCC) and the cumulative profiles of n=23 PCAWG TCC (top). The difference (removing negative contributions) compared to the SBS-MM1 mutational signature profile (bottom).

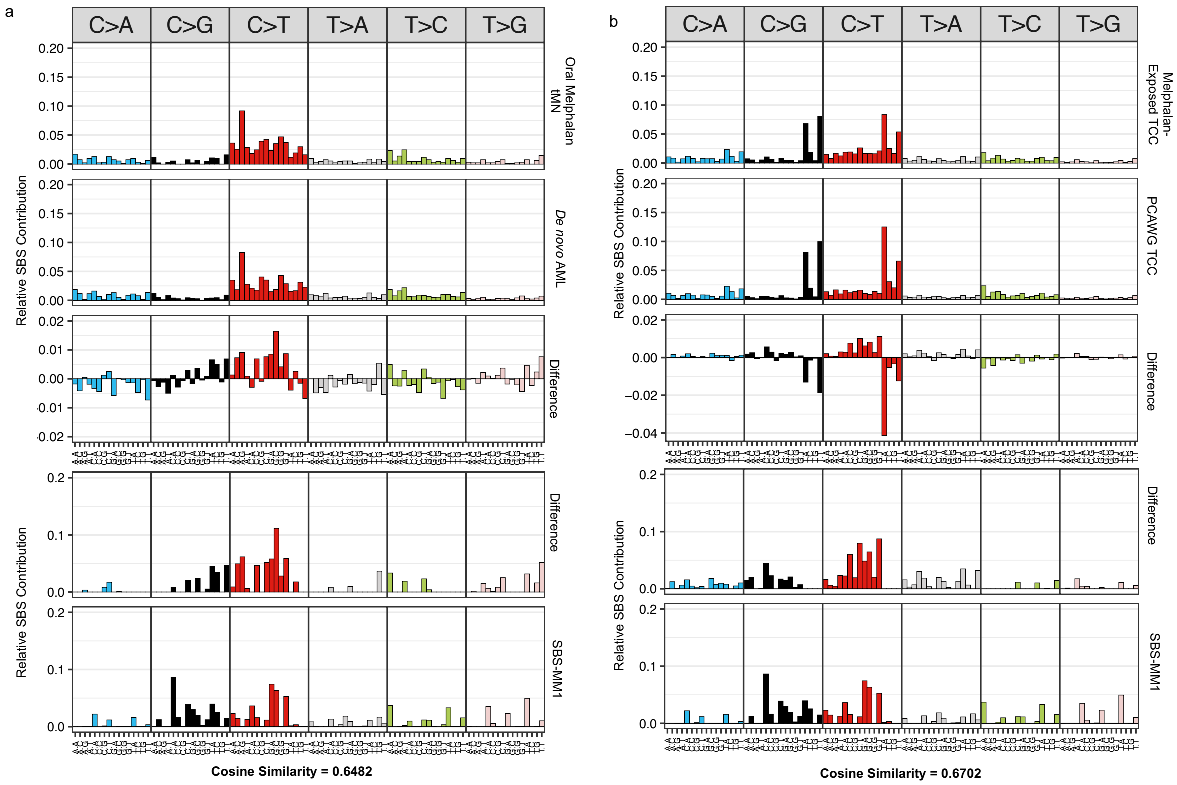

**Extended Data Figure 8. Mutations in driver genes in therapy related myeloid neoplasms. a)** Oncoplot of driver SNV defined using *dndscv* for 316 *de novo* AML and 61 tMN. **b)** Pie plots of SBS signature contribution to pooled mutations in driver genes from genomes of *de novo* AML (top, 25 SNV), chemotherapy-signature-negative tMN (middle, 31 SNV), and chemotherapy-signature-positive tMN (bottom, 32 SNV).

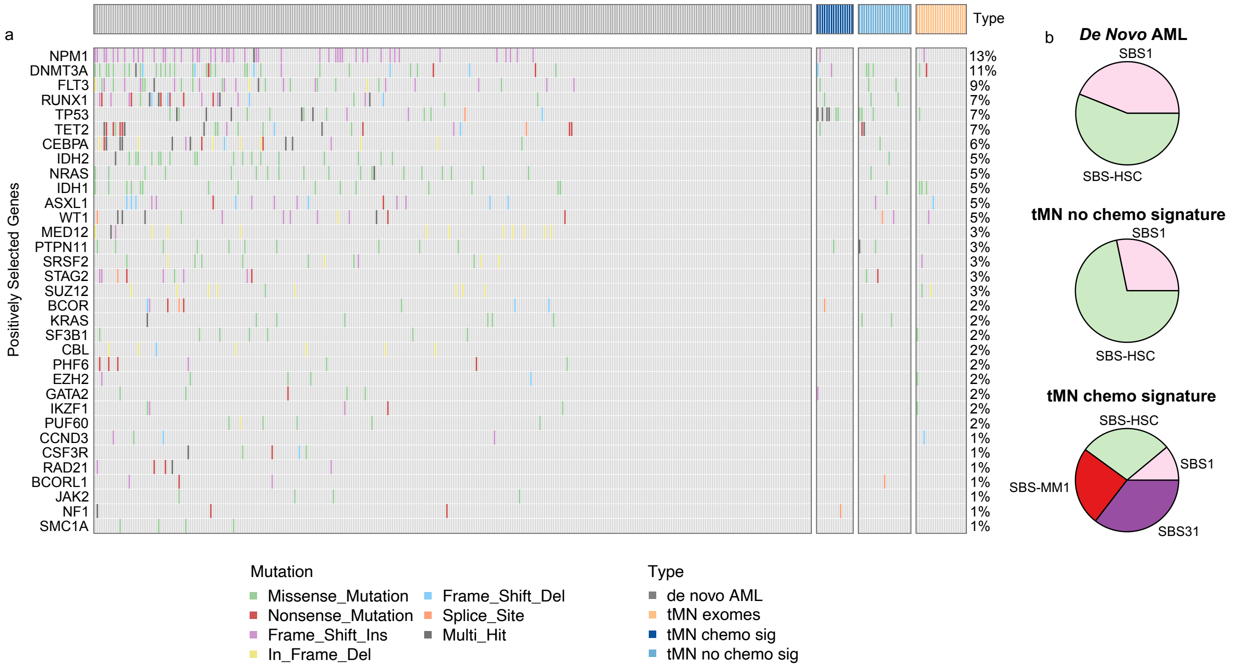

**Extended Data Figure 9. Copy number and structural variant differences in therapy-related myeloid neoplasms with and without chemotherapy-associated single base substitution signatures. a)** Cumulative copy number profile or tMN genomes with a chemotherapy mutational signature (n = 17, top), tMN genomes without a chemotherapy signature (n=22, middle), and tMN exomes from Beat AML (n= 22). **b)** Structural variant landscape across tMN genomes with (top) and without (bottom) a chemotherapy-associated mutational signature. Breakpoints are binned into 1 megabase segments. Simple events point upward from the x-axis and complex events point downward. **c)** Boxplot of number of complex structural variants by presence of chemotherapy signature among tMN cases. The p-value was estimated using Wilcoxon test.

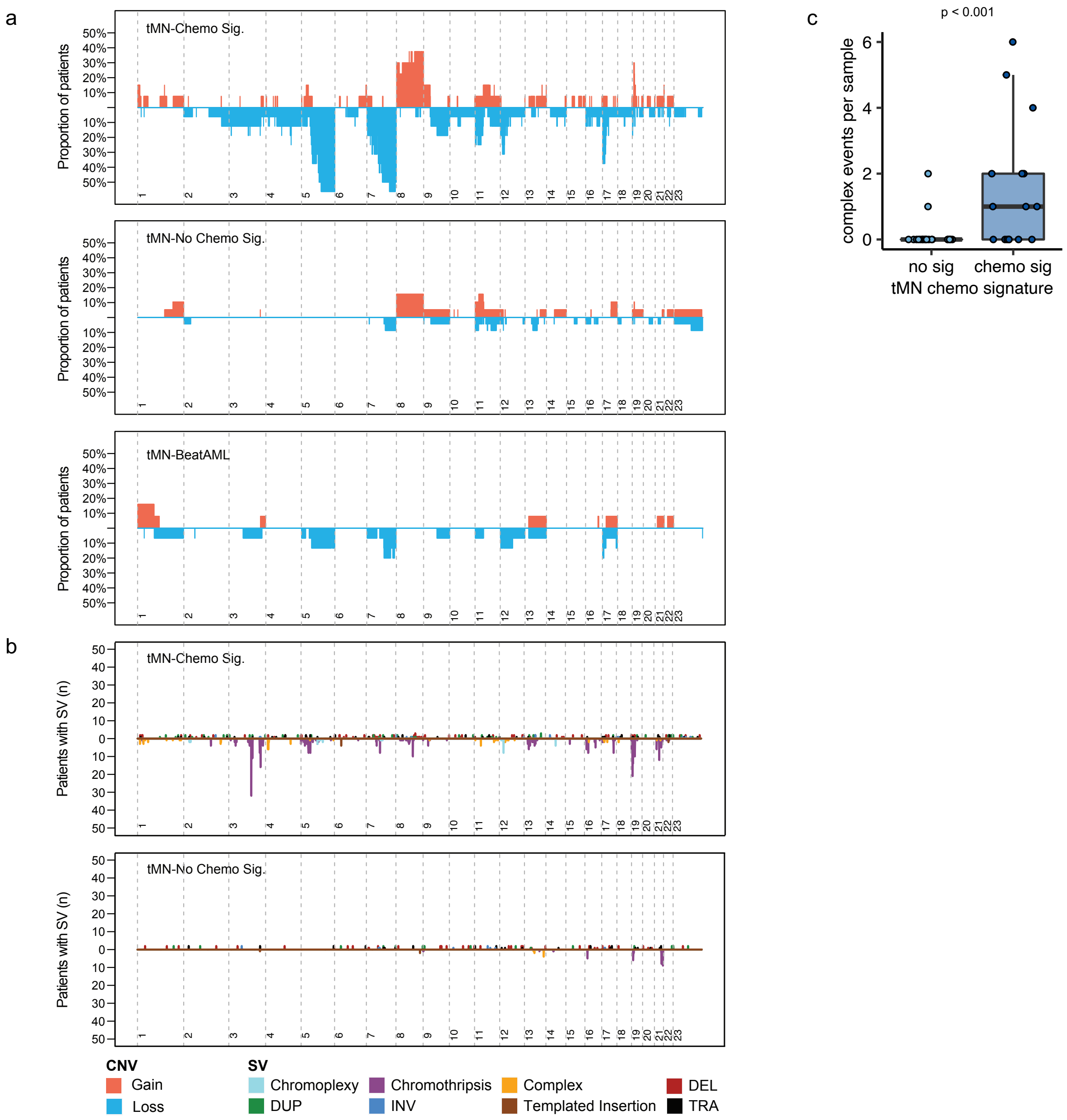

**Extended Data Figure 10. Simple and complex structural variants involving the *SMARCA4* and *MECOM* loci in tMN genomes.** The horizontal black line indicates the total copy number; the dashed orange line indicates the minor copy number. The vertical lines represent SV breakpoints, color-coded based on SV class: blue = inversion, green = tandem-duplication; red = deletion; black = translocation.

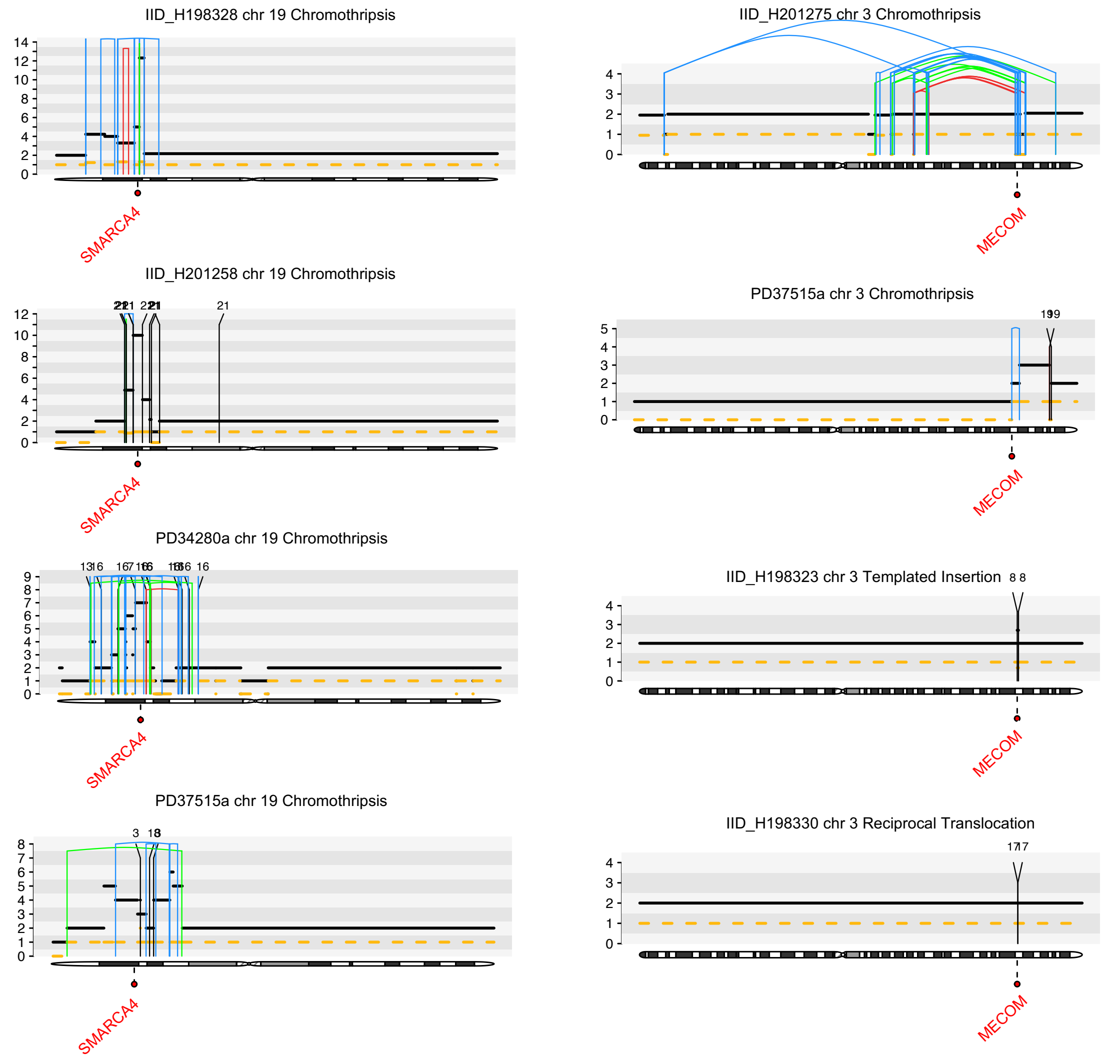

**Extended Data Figure 11. *SMARCA4* in *de novo* AML and Ba/F3 cell line. a)** Copy number profile from the only *de novo* AML tumor (among n=298 cases in the BEAT-AML study) to contain a high copies amplification of the *SMARCA4* locus. **b)**: Western blot showing expression of *SMARCA4* in transfected cells compared to vector for cytokine independence assay.

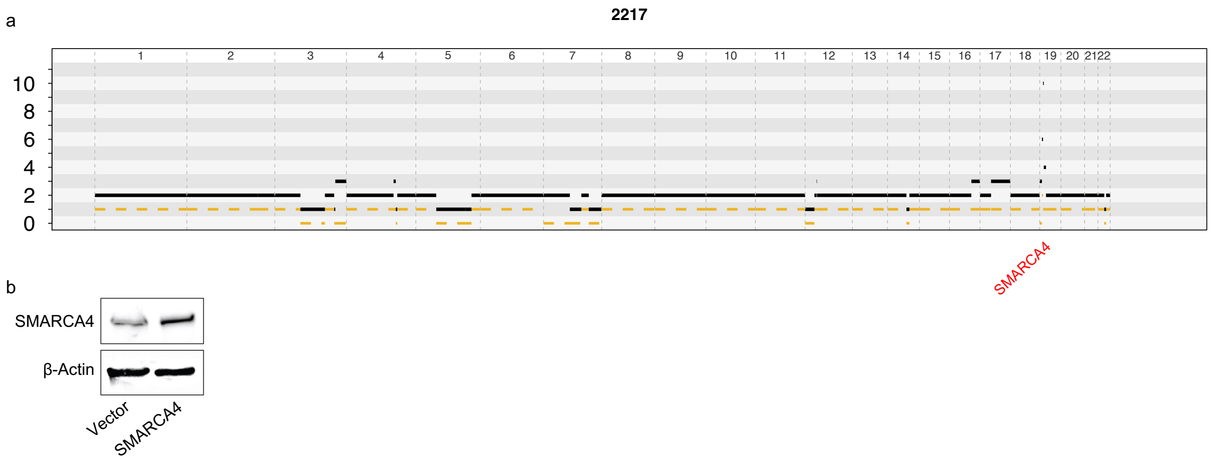

**Extended Data Figure 12. Molecular time and clock-like single base substitution mutation rate in multiple myeloma, *de novo* and therapy-related myeloid neoplasms. a)** Linear regressions showing the (lack of) association between clock-like SBS mutations and age of tumor, regardless of presence or absence of chemotherapy-induced mutagenesis (i.e., chemotherapy-associated mutational signatures). P-values and R squares were estimated using *lm* R function. **b)** Molecular time estimates for large gains in eligible tumors. Events occurring closer to tumor sequencing (i.e., diagnosis) are later in molecular time (i.e., closer to 1). **c)** Individual SBS5 mutation rate estimate for multiple myeloma using linear mixed effect model. The SBS5 mutational burden was derived from the phylogenetic branches of each patient (dots). A total of 77 WGS (multiple myeloma and smoldering myeloma) from 47 patients cases from a prior study [8] were included together with two newly sequenced post-platinum multiple myeloma tumors.

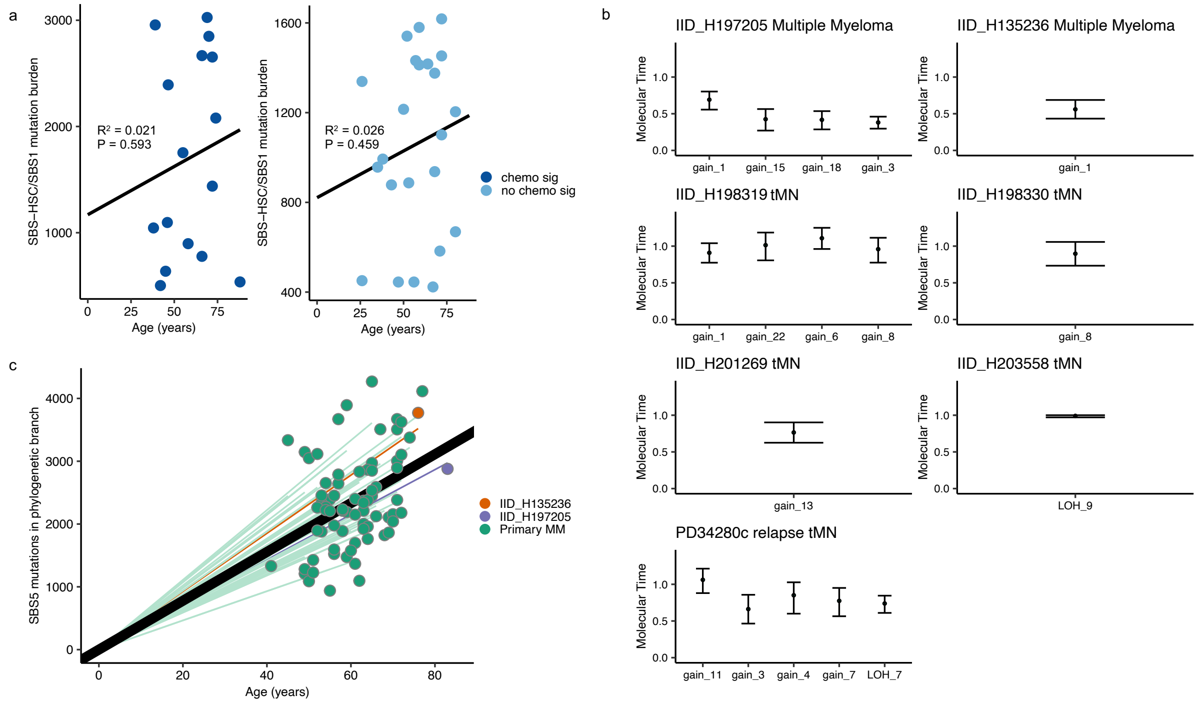

**Extended Data Figure 13. Example of Clonal Hematopoiesis.** Screenshot from Integrated Genome Viewer (IGV) showing a small *TP53* mutant clone in a sample taken from the leukapheresis product. This clone expanded into a tumor without a melphalan signature, having escaped direct exposure to chemotherapy and resultant mutagenesis.

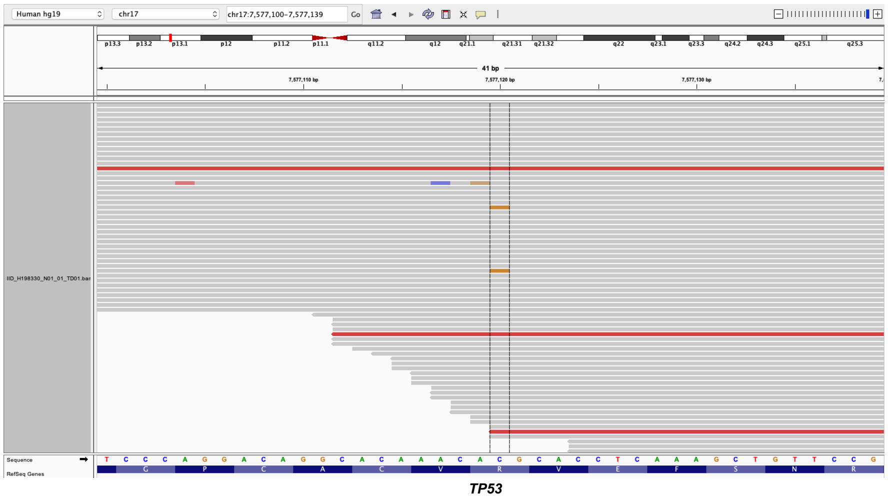
